# Coordinated drift of receptive fields during noisy representation learning

**DOI:** 10.1101/2021.08.30.458264

**Authors:** Shanshan Qin, Shiva Farashahi, David Lipshutz, Anirvan M. Sengupta, Dmitri B. Chklovskii, Cengiz Pehlevan

## Abstract

Long-term memories and learned behavior are conventionally associated with stable neuronal representations. However, recent experiments showed that neural population codes in many brain areas continuously change even when animals have fully learned and stably perform their tasks. This representational “drift” naturally leads to questions about its causes, dynamics, and functions. Here, we explore the hypothesis that neural representations optimize a representational objective with a degenerate solution space, and noisy synaptic updates drive the network to explore this (near-)optimal space causing representational drift. We illustrate this idea in simple, biologically plausible Hebbian/anti-Hebbian network models of representation learning, which optimize similarity matching objectives, and, when neural outputs are constrained to be nonnegative, learn localized receptive fields (RFs) that tile the stimulus manifold. We find that the drifting RFs of individual neurons can be characterized by a coordinated random walk, with the effective diffusion constants depending on various parameters such as learning rate, noise amplitude, and input statistics. Despite such drift, the representational similarity of population codes is stable over time. Our model recapitulates recent experimental observations in hippocampus and posterior parietal cortex, and makes testable predictions that can be probed in future experiments.

## Introduction

Memories and learned behavior can be stable for a long time. We can recall vividly the memory of events that happened years ago. Motor skills, such as riding a bike, once learned, can last life-long even without further practice. A natural question is then whether stable task performance and memories are related to stable neuronal representations.

Recent technical advances in electrophysiology and optical imaging enabled researchers to address this question by studying the long-term dynamics of neural population activity in awake behaving animals (Katlowitz, Picardo, and Long 2018; Li et al. 2017; Luo et al. 2020; Schoonover et al. 2021; Ulivi et al. 2019; Y. Ziv et al. 2013). A number of these experiments have shown that neuronal activities in cortical areas that are essential for specific tasks undergo continuous reorganization even after the animals have fully learned their tasks, a phenomenon termed “representational drift” (Mau, Hasselmo, and Cai 2020; Rule, O’Leary, and Harvey 2019). For instance, in sensorimotor tasks, neuronal representations in the posterior parietal cortex (PPC) of mice change across days while the performance of animals remain stable and high (Driscoll et al. 2017). Place fields of individual place cells in CA1 region of hippocampus drift over days and weeks even when the animals remain in the same familiar environment (Gonzalez et al. 2019; Lee et al. 2020; Y. Ziv et al. 2013). Individual neurons in the primary motor cortex and supplementary motor cortex show unstable tuning while animals perform highly stereotyped motor tasks (Rokni et al. 2007) (but see (Chestek et al. 2007; Gallego et al. 2020)). Representational drift has been observed even in primary sensory cortices, such as mouse visual cortex (Deitch, Rubin, and Y. Ziv 2021; Marks and Goard 2021) and piriform cortex (Schoonover et al. 2021). The ubiquity of representational drift raises several important questions: What is the underlying mechanism of such drift? How can neural circuits generate stable coding in the presence of continuous drift? What are the dynamics of representational drift?

To address these issues, we consider a setting where a neural population learns to represent stimuli in a way that optimizes a representational objective. Such a normative account of sensory representations is common in neuroscience (Atick and Redlich 1992; Attneave 1954; Barlow 1961; Chalk, Marre, and Tkacik 2018; Hateren 1992; Olshausen and Field 1997; Pehlevan, Hu, and Chklovskii 2015; Rao and Ballard 1999; Srinivasan, Laughlin, and Dubs 1982). Furthermore, the representational objective we consider has many solutions, consistent with the notion that there are many optimal neural representations of input stimuli. Based on the observations that synapses in the cortex are highly dynamic (Attardo, Fitzgerald, and Schnitzer 2015; Hazan and N. E. Ziv 2020; Rumpel and Triesch 2016), we hypothesize that noisy synaptic updates during learning will drive the network to explore the synaptic weight space that corresponds to (near-)optimal neural representations. In other words, the neural representation will drift within the space of optimal representations.

To test this hypothesis, we focus on a well-studied biologically plausible network for representation learning: the Hebbian/anti-Hebbian network (Földiak 1990; Pehlevan and Chklovskii 2019) (Fig. 1B). These networks optimize similarity matching objectives which exhibit a degeneracy of optimal representational solutions (Pehlevan, Sengupta, and Chklovskii 2018), all of which share the same representational similarity matrix (Kriegeskorte, Mur, and Bandettini 2008). Hebbian/anti-Hebbian networks have also been shown to learn localized RFs that tile the input data manifold (Sengupta et al. 2018), hence they can be used as simplified models for brain areas where neurons have localized receptive fields (RFs), such as hippocampal place cells and neurons in PPC (Driscoll et al. 2017; Gonzalez et al. 2019; Y. Ziv et al. 2013). In these systems, population of neurons with different localized RFs generally tile the parameter space they encode (Fig. 1A). The simplicity and mathematical-tractability make Hebbian/anti-Hebbian networks an excellent starting point to elucidate the mechanism and properties of representational drift due to noise in synaptic updates.

**Figure 1:**
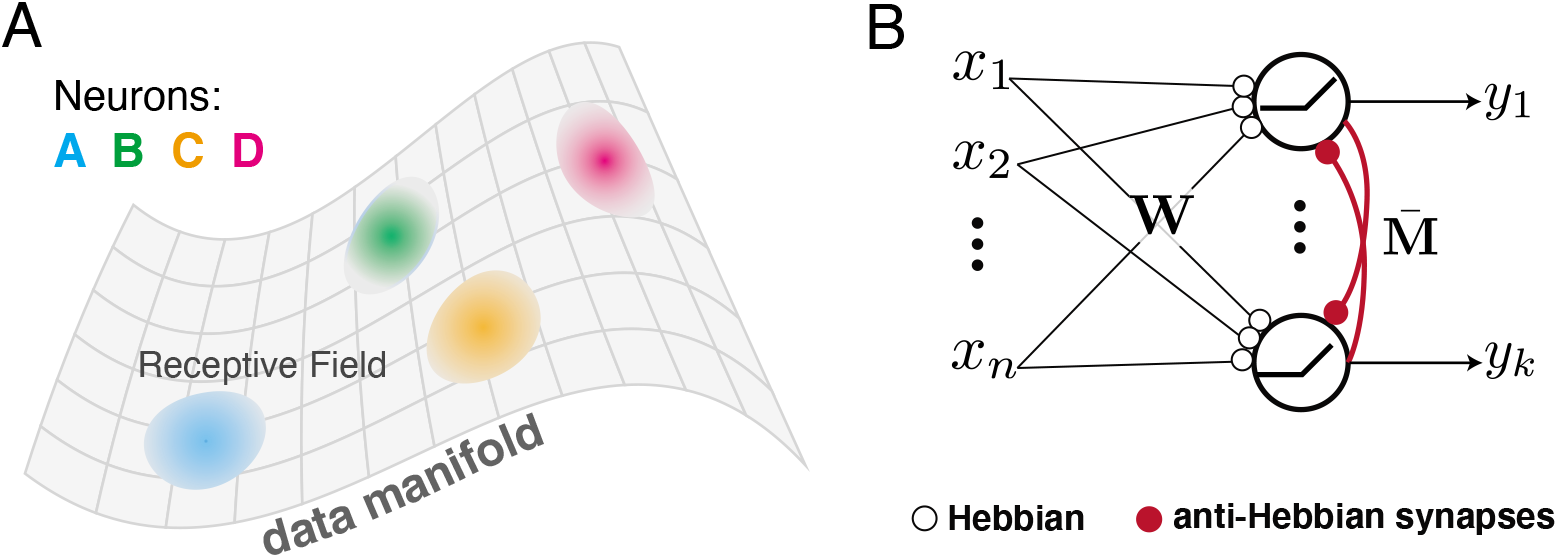
(A) Illustration of localized receptive fields that tile the data manifold. (B) Hebbian/anti-Hebbian network with nonnegative neural activity can learn localized receptive fields.

By numerical and analytical methods, we find that while the RFs of individual neurons change significantly over time, the representational similarity of population codes is stable.We show that the drift dynamics of individual RFs can be largely captured by a random walk on the data manifold, with the effective diffusion constant depending on noise amplitude, learning rate and other model parameters such as the number of output neurons. At the population level, the drifting RFs are coordinated in a way that preserves representational similarity. Our model accounts for many of the recent experimental observations in the hippocampus and PPC, and makes testable predictions. Overall, our results show how optimal representation learning and noise can lead to representational drift while maintaining representational similarity.

## Results

### Drift dynamics in linear Hebbian/anti-Hebbian networks

We first study drift in linear Hebbian/anti-Hebbian networks, which compress inputs into a lower dimensional space (Pehlevan, Hu, and Chklovskii 2015). While the resulting RFs are not localized, it is still instructive to study how learned representations evolve with noisy synaptic updates in this analytically tractable model.

The network we consider minimizes a similarity matching cost function (Pehlevan, Hu, and Chklovskii 2015). Here, the similarity between two vectors is defined as their dot product. Let **x***_t_* ∈ ℝ^*n*^, *t* = 1, …, *T* be a set of network inputs (or sensory stimulus) and **y***_t_* ∈ ℝ^*k*^, *k* < *n* be the corresponding outputs constituting a representation. Similarity matching minimizes the mismatch between the similarity of pairs of inputs and corresponding pairs of outputs

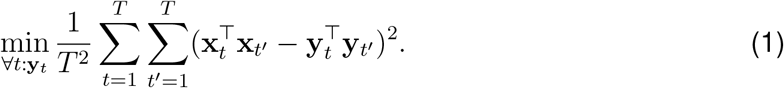

Optimal solutions to this problem are given by projecting the inputs to their principal subspace (Pehlevan, Hu, and Chklovskii 2015). However, there is a continuum of such projections each corresponding to a basis in the subspace. This degeneracy can be seen from the rotational symmetry of the similarity matching cost function, (1). For any set of **y***_t_*, **Ry***_t_* has the same cost, where **R** is an orthogonal matrix.

Previous work showed that this cost function can be minimized by a neural network in an online manner, where each input **x***_t_* is presented sequentially and an output **y***_t_* is produced (Pehlevan, Hu, and Chklovskii 2015) (Materials and Methods) by running the following neural dynamics until convergence:

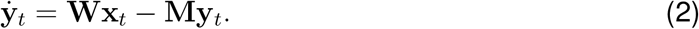

Here, **W** holds the feedforward synaptic weights and **M** the lateral weights. We note that at the fixed point of the neural dynamics **y***_t_* = **M**^−1^**Wx***_t_* = **Fx***_t_*, where we define a filter matrix, **F** ≡ **M**^−1^**W**, whose rows are neural filters. After each presentation of an input and convergence of the neural dynamics, the weights **W** and **M** are updated with a Hebbian and anti-Hebbian rule, respectively:

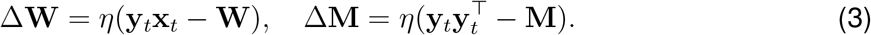

The learning rule is local in the sense that synaptic update only depends on activities of presynaptic and postsynaptic neurons. The update of **M** is anti-Hebbian due to the negation in (2). As the number of samples increases, these weights converge to a configuration where neural filters learn an orthonormal basis for the principal space (Pehlevan, Hu, and Chklovskii 2015; Pehlevan, Sengupta, and Chklovskii 2018).

Here, we model biological noise during learning by introducing noise to the weight updates, and examine the consequences of this noise. The updates are

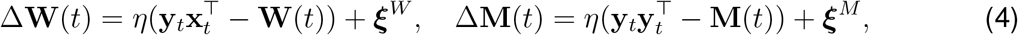

where the noise terms 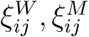 are independent Gaussian noise, with the following statistics: 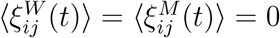 and 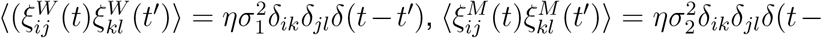 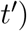.

As expected, the network learns the principle subspace and maintains a stable performance, as quantified by the principle subspace projection (PSP) error: 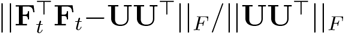 where **U** is a *n* × *k* matrix whose columns are the top *k* left singular vector of **X** (Fig. 2A). Due to the synaptic noise, network weights do not settle down to fixed points but roam around in the subspace that gives equally good solutions to the similarity matching problem. Consequently, the representation of a given stimulus **y***_t_* drifts over time (Fig. 2B). However, the similarity between any two outputs **y***_t_* and **y***_t′_* remains stable (Fig. 2C, SI Appendix Fig. S1A). As a consequence, the drift does not change the length of the output vectors, **y***_t_*, which undergo a random walk on a spherical surface (Fig. 2D). The drift of the representation ensemble **Y***_t_* ≡ [**y**_1_, …, **y***_T_*] behaves like a randomly-rotating rigid body consisting of a cloud of points (SI Appendix Fig. S1B).

**Figure 2:**
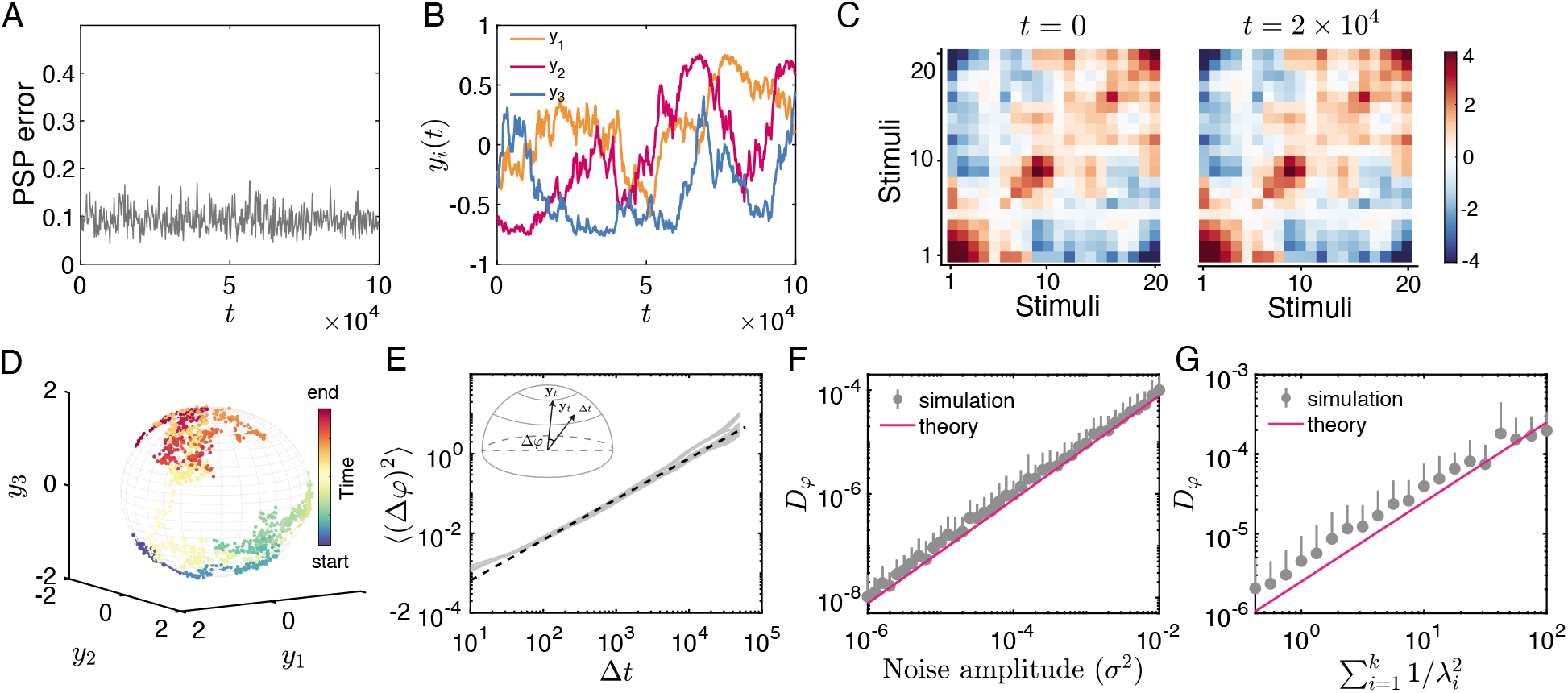
Drift dynamics in principal subspace projection (PSP) task. In the simulation, each input **x**_*t*_ ∈ **R**^10^ is drawn independently from a joint Gaussian distribution 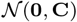. The first three eigenvalues of the covariance matrix **C** are: 4.5, 3.5, 1 and the rest are 0.1. A Hebbian/anti-Hebbian network learns to project this input to *k* = 3 dimensions. (A) PSP error remains stable after the task has been learned even with noisy synapse update. (B) The learned representation of an example input continuously changes due to noisy updates. Shown are the 3 component of **y**(*t*). (C) Pairwise similarity between learned representations are stable over time, as shown by the almost identical similarity matrices at *t* = 0 (left) and *t* = 2 × 10^4^ (right). (D) Drifting representation as a random walk on a “sphere”, showing the representation of a single sample **y***t* over time. Color codes for different time steps. (E) Estimating rotational diffusion constant *D_φ_* from mean squared angular displacement (MSAD). Grey lines are MSAD estimated based on individual representation trajectory **y**(*t*). The dashed line is a linear fit between 〈(Δ*φ*)^2^〉 ≡ 〈(*φ*(*t* + Δ*t*) − *φ*(*t*))^2^〉 and Δ*t* to estimate the rotational diffusion constant. Inset: illustration of Δ*φ*. (F) Relationship between *D_φ_* and noise amplitude *σ*^2^. Symbol with error bars are numerical simulation, and the solid line is the theory Eq.5. (G) Dependence of *D_φ_* on the eigenspectrum {*λ_i_*} of the input covariance matrix. Error bars represent standard deviation, only one side is shown to reduce cluttering. In all the figures *t* = 0 is the starting point when the representation is learned. Parameters: *n* = 10, *k* = 3, *η* = 0.1, *σ*_1_ = *σ*_2_ = 0.01, *T* = 10^4^.

These observations motivated us to quantify the drift rate by the rotational diffusion constant *D_φ_* (Kämmerer, Kob, and Schilling 1997; Mazza et al. 2006) (Materials and Methods). We can derive an approximate analytical formula for *D_φ_* from a linear stability analysis of the filter matrix (**F**) (Materials and Methods, and SI Appendix for details):

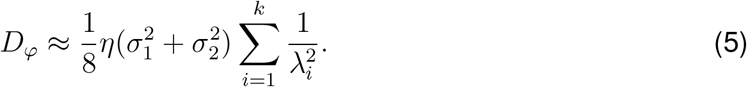

Here, *λ*_1_ ≥ *λ*_2_ ≥ … ≥ *λ_k_* are the ordered eigenvalues of the input covariance matrix. (5) indicates that *D_φ_* is proportional to the noise amplitude (Fig. 2F). Further, the drift amplitude along each eigenvector is proportional to 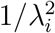 This is analogous to the rotation of an ellipsoid rigid body due to torque. In that system, the rotation around the axis with smaller moment of inertia is easier. Predictions of (5) is well in agreement with simulations (Fig. 2E-G).

The above simulation and analysis demonstrate that, in this model, while the network’s output is drifting over time, similarity of representation is preserved. This is due to a coordinated random walk in the representational space, which in the linear case can be described by a rigid body rotation. This coordinated drift explores equally optimal representations. Because the quality of a representation is quantified by its representational similarity in (1), representational similarity is preserved despite the drift. Next, we consider nonlinear networks and show that these results carry over.

### Drift dynamics in nonlinear Hebbian/anti-Hebbian networks

RFs of neurons in many brain areas are localized in the parameter space which they represent. For example, response of neurons in primary visual cortex (V1) are tuned to orientations of gratings (Hubel 1995). Neurons in the owl’s external nucleus of the inferior colliculus (ICX) are tuned to different horizontal and vertical positions, forming an auditory spatial map (Peña and Konishi 2001). Place cells in hippocampus are active when an animal is at a particular spatial location of an environment (O’Keefe and Dostrovsky 1971).

A nonlinear version of our Hebbian/anti-Hebbian network can capture these localized RF properties (Fig. 1B). This network minimizes a nonnegative similarity matching (NSM) cost function (Földiak 1990; Pehlevan, Hu, and Chklovskii 2015; Sengupta et al. 2018):

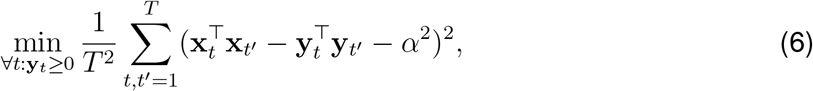

where **x***_t_* ∈ ℝ^*n*^ and **y***_t_* ∈ ℝ^*k*^ are input and output, respectively, and *α*^2^ sets the threshold of similarity to be preserved in the output representation. With nonnegative neural activity, the above NSM objective function strives to preserve similarity for similar pairs of input samples but orthogonalizes the outputs corresponding to dissimilar input pairs. Compared with the linear network, non-negativity breaks the rotational symmetry of the solution, but the permutation symmetry is still preserved, i.e., exchanging identities of neurons does not change the objective function. To see this more clearly, the above target function can be written in terms of input-output Gram matrices: 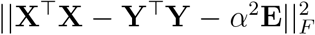, where **X** ∈ ℝ^*n*×*T*^, **Y** ∈ ℝ^*k*×*T*^, and **E** ∈ ℝ^*T* ×*T*^ is a matrix with all entries set to 1. Thus, if **Y** is a solution, then **PY** is also a solution for any permutation matrix **P**. Note that a general rotation would not preserve the nonnegativity of the output and thus not lead to a new solution. One can further control the behavior of learned representations by introducing regularizers to **y***_t_* in (6), for example an *l*_1_ norm of **y** leads to more sparse code (Materials and Methods).

Similar to the previous linear Hebbian/anti-Hebbian network, this network also operates in an online fashion with a similar local learning rule. It takes an input **x***_t_* and generates an output **y***_t_* by running the following neural dynamics until it converges (Pehlevan 2019):

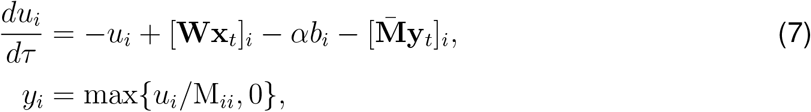

where *u_i_* and *y_i_* represent the membrane potential and firing rate of neuron *i*, and *b _i_* is the bias term. The forward weight matrix **W** ∈ ℝ^*k*×*n*^ and lateral weight matrix **M** ∈ ℝ^*k*×*k*^ (we have defined 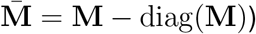 update according to the following “noisy” learning rule (Pehlevan 2019):

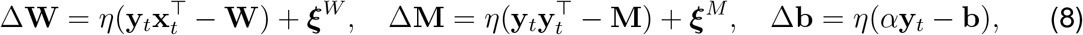

where *η* is the learning rate, and ***ξ****^W^* and ***ξ****^M^* are Gaussian white noise terms with the same statistics as in (4). The properties of the above learning rule without noise has been studied previously (Földiak 1990; Pehlevan 2019; Pehlevan and Chklovskii 2014; Pehlevan, Mohan, and Chklovskii 2017; Sengupta et al. 2018). Here, our interest is investigating how the learned representations evolve in the presence of noise in synaptic updates. We will first study the general drifting dynamics of RFs by a simple input: a ring data manifold. Based on the insights gained from this model, we will build models of drifting representations in place cells and neurons in PPC.

### Localized receptive fields on a ring stimulus manifold

To explore how RFs evolve in the presence of synaptic noise, we first consider stimuli living on a ring (Fig. 3A), like the direction of a drifting grating used in experimental study of visual systems. Location of the stimulus on the ring is parameterized by an angular variable *θ* ∈ [0, 2*π*) (Materials and Methods). The input similarity matrix **X**^┬^**X** has a diagonal band structure and two inputs that are close on the ring have large similarity.

**Figure 3:**
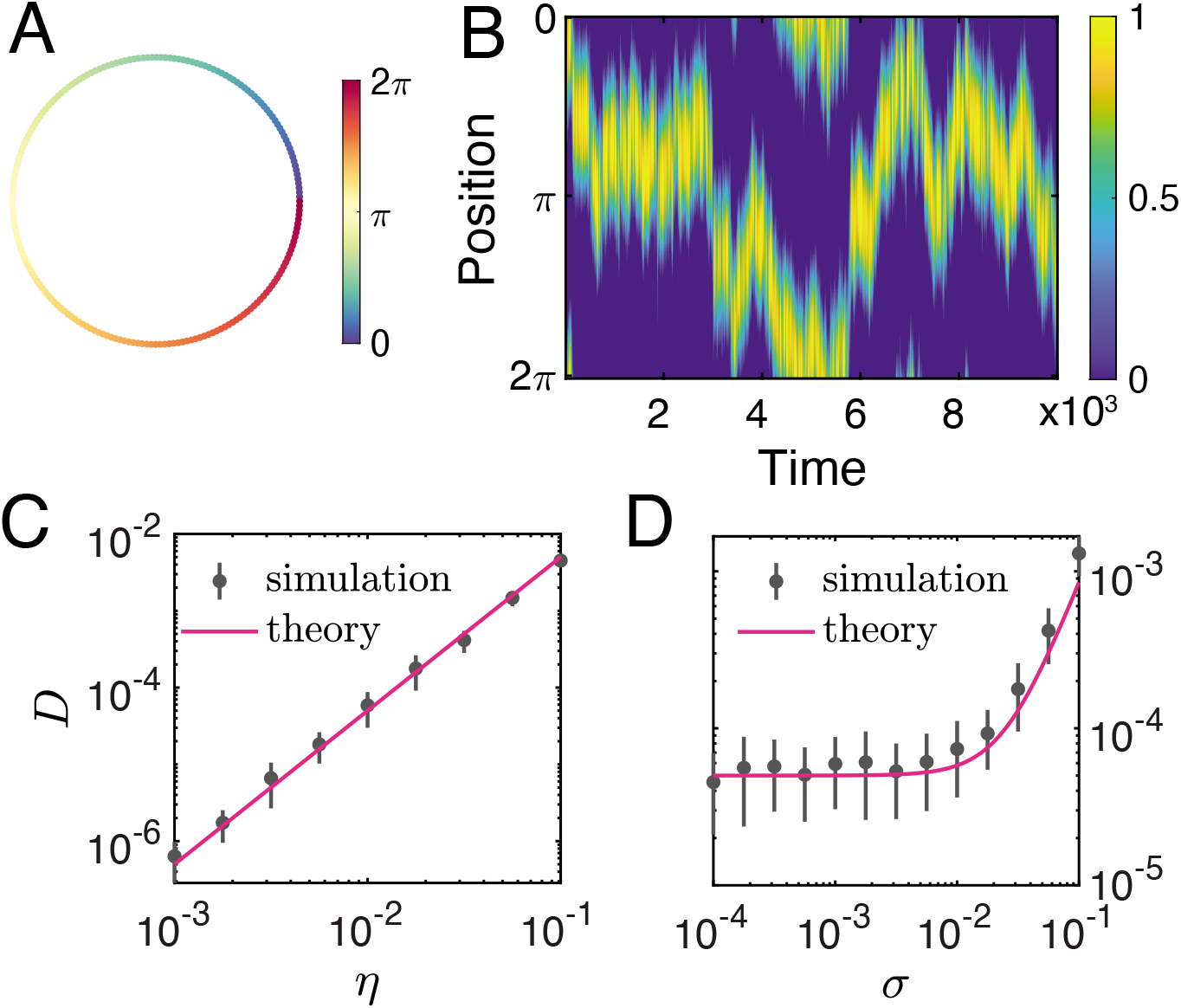
Drift of a single localized RF learned from a ring data manifold. (A) A ring in 2D as input dataset: **x**(*θ*) = [cos(*θ*), sin(*θ*)]^┬^, *θ* ∈ [0, 2*π*). (B) The single RF has the shape of a truncated cosine curve, whose position drift on the ring and behaves like a random walk. (C,D) The effective diffusion constant *D* of centroid position increases with learning rate *η* even without explicit synatpic noise (*σ* = 0), and with the noise amplitude of explicit synaptic noise. Error bars represent standard deviation of 40 simulations. Megenta lines correspond to (9). Parameters: *α*^2^ = 0, *β*_1_ = *β*_2_ = 0, in (D) *η* = 0.05.

In the case of a single output neuron and in the absence of noise, the learned RF can be solved analytically, which is a truncated cosine curve centered around a random angle *ϕ*, i.e., *y_ϕ_*(*θ*) = *μ*[cos(*θ* − *ϕ*)]_+_ with *μ* being the peak amplitude. Derivation of this results and dependence of *μ* to model parameters is given in Materials and Methods. With synaptic noise during learning, the centroid drifts on the ring like a random walk. We quantified the speed of drift with an effective diffusion constant, *D*, of the centroid by the conventional relation: 〈(*ϕ*(*t* + Δ*t*) − *ϕ*(*t*))^2^〉 = 2*D*Δ*t*, where *ϕ*(*t*) is the centroid position at time *t*. For a single neuron, the dependence of *D* on the learning rate *η* and noise amplitude *σ*^2^ can be analytically approximated as (Materials and Methods, and SI Appendix):

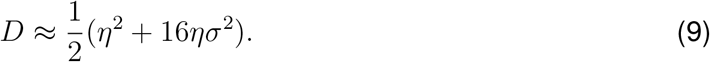

The first term of (9) is due to the sampling noise, i.e., the fact that the network sees one random stimulus at a time, and the second term is due to the explicit synaptic noise (noise terms in (8)). (9) indicates that faster learning and larger explicit noise result in more rapid drift of the RF. Numerical simulation agrees well with the theory (Fig. 3C-D).

When there is a population of output neurons, the Hebbian/anti-Hebbian network learns multiple localized RFs that tile the ring manifold with overlap (Fig. 4A), consistent with previous analytical accounts of simplified versions of such networks (Sengupta et al. 2018). With synaptic noise, we expect each RF to drift by a similar diffusion process as in the single neuron case, but with interactions between the neurons affecting the dynamics (Fig. 4B). In particular, the structure of neural population activity, as indicated by output similarity matrix **Y**^┬^**Y**, is stable across time (Fig. 4C). Further, a neuron’s response to the same stimulus is intermittent, having active and silent periods (Fig. 4D). At the population level, the fraction of neurons that have active RFs at any given time is constant (Fig. 4E), and it decreases with total number of output neurons as well as the noise amplitude (Fig. 4F). Thus, in a large population of neurons, only a small fraction of them will be active at a give time, forming a sparse population code. Neurons whose RFs have stronger tuning (characterized by the peak amplitude of RF) tend to be active more often (Fig. 4G), and have smaller drift (Fig. 4H).

**Figure 4:**
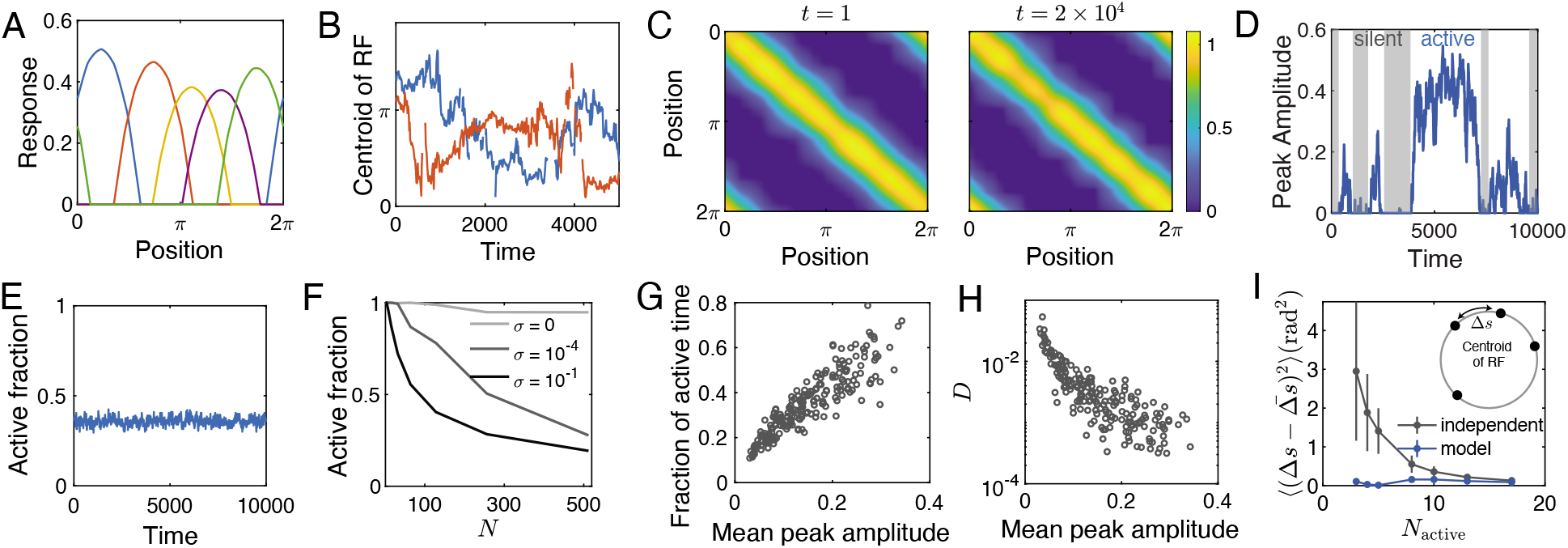
Drift of manifold-tiling localized RFs. (A) Learned localized RFs tile the input ring data manifold. Colors represent RFs of 5 example neurons. (B) Evolution of the RF centroids of two example neurons due to synaptic noise. (C) The representational similarity matrix **Y^┬^Y** is approximately circulant and stable over time. (D) When there are large number of neurons, each neuron has active and silent (shaded region) periods. (E) At population level, the fraction of neurons with active RFs are constant. (F) The fraction of neurons that have active RFs decreases with the total number of output neurons, as well as the noise amplitude. (G-H) Neurons that have stronger RFs tend to have longer active time (G) and also are more stable as quantified by smaller effective diffusion constants *D* (H). (I) At population level, the drift of RFs are coordinated such that their centroids are more uniformly distributed compared to that of the independent random walk case, in which the step size follows the same distribution of the Hebbian/anti-Hebbian network model. Shown are the variance of distances between adjacent centroids. Parameters in C-G: *N* = 200, *σ* = 0.002, *η* = 0.05, *α*^2^ = 0 except *σ* = 0 in B. In H: *η* = 0.05.

At the population level, the drift of RFs are coordinated. To see the difference between our model and independent random walkers, we simulated *N*_active_ centroids undergoing independent random walks on the ring. The step size of the independent random walks were drawn from the same distribution as the centroid shift between two adjacent time steps in our model (Materials and Methods). We observed that centroids in our model tile the ring manifold more uniformly than those of independent random walks, as indicated by the smaller variance of distances between two adjacent centroids on the ring (Fig. 4I).

Having gained better understanding of drifting dynamics in the above simple model, we now discuss models of representational drift in the Hippocampus CA1 region and PPC. The observations made in Fig. 4 will conceptually carry over, providing explanations for previous experimental observations.

### A Hebbian/anti-Hebbian Network model for drifting place fields in CA1

CA1 place cells in the hippocampus play a crucial role in spatial memory and navigation. Recent long-term recording experiments show that place coding by the population of CA1 pyramidal cells are dynamic even when the animal is in the same familiar environment. In the time course of several weeks, some neurons lose their place fields while other previously non-place coding cells gain place fields. Despite the drift, the spatial information is preserved (Gonzalez et al. 2019; Y. Ziv et al. 2013).

One possible mechanism of place field formation is that CA1 place cells receive both forward input from grid cells in the entorhinal cortex and lateral competition from other place cells within the hippocampus (M.-B. Moser, Rowland, and E. I. Moser 2015). This motivated us to use the Hebbian/anti-Hebbian network to learn a place cell representation of a 2D square environment. Each position on the plane is represented by a population of grid cells with different grid spacing, phases and offsets (Fig. 5A, Materials and Methods), which serves as the input **x***_t_* to the network. After learning, some output neurons develop localized RFs (or place fields, Fig. 5B). This can be visualized by arranging each row of response matrix **Y** into a square matrix, as shown in Fig. 5B. The population of output neurons tile the 2D environment, as indicated by the uniform distribution of centroids of place fields (Fig. 5B, right). Due to the noise in the weight update, place fields continuously drift over time (Fig. 5C). Despite the drift, representational similarity of positions in the 2D environment is stable (SI Appendix, Fig. S2A). We also observed that a place cell may lose its place field for some time and gain a new place field later on (SI Appendix, Fig. S2B). The intermittence of RFs is due to both the competition between RFs and the fluctuation of **W** and **M**. For example, once the forward input is smaller than the lateral inhibition at the centroid of an RF, then this RF becomes silent. The interval of these silent periods follow exponential distributions (SI Appendix Fig. S2C), indicating that the transition between active and silent state of RFs are random and memoryless. However, the fraction of neurons with active place fields at any given time remains constant (SI Appendix, Fig. S2D).

**Figure 5:**
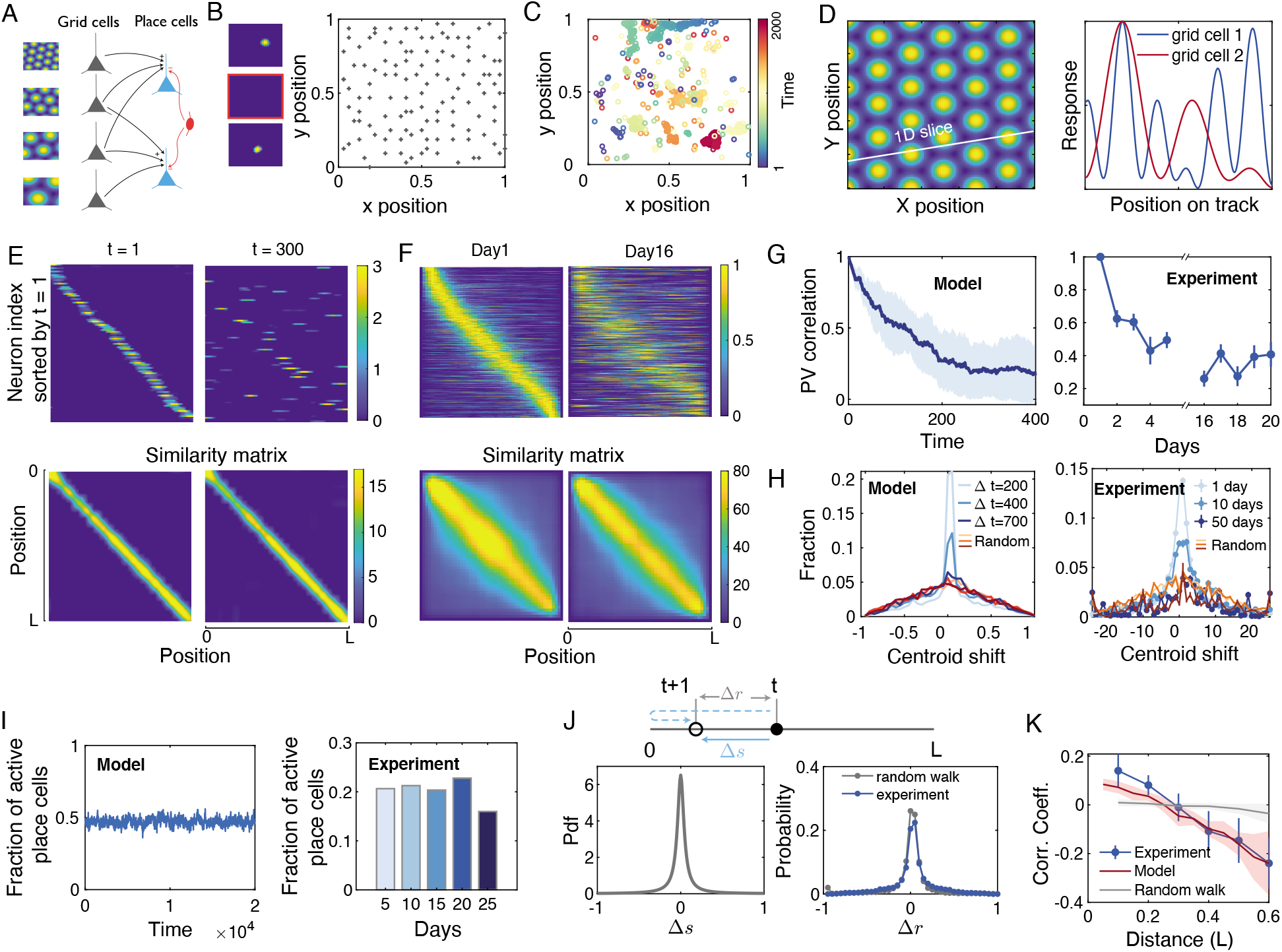
Drift of place fields. (A) Place cells receive input from grid cells that have different grid spacing, orientations and offsets. They also receive lateral inhibition due to competition with other place cells. (B) Left: learned place fields (left) tile the entitle 2D plane with red square highlighting an silent neuron. Right: each dot represents a centroid of a place field. (C) Drift of place field of an exemplar place cell. Each circle represents the position of its centroid at different times. (D) Left: Slice through a 2D grid field. Right: Response of neurons across this slice. (E) Upper: learned place fields tile a 1D linear track when sorted by their centroid positions (left), but continuously change over time (right). Lower: Representational similarity matrix **Y^┬^Y** of position is stable over time. (F) Experimental results corresponding to (E), place fields of a group of CA1 place cells concatenated from several mice when exploring the same familiar 1D linear track. (G) The average autocorrelation coefficient of population vectors representing each spatial position in the model (left) and experiment (right) decay over time. Shades represents standard deviation over different positions. (H) Probability distribution of centroid drifts of place cells at three different time intervals. Red lines represent random distributions, which are obtained by randomly aligning place fields of neurons between the same interval. The qualitative behavior of the model (left) is very similar to that of the experimental result (right). (I) Despite the continuous reconfiguration of place cell ensembles, the fraction of active place fields are stable over time in the model (left) and experiment (right). (J) Centroid shift Δ*r* = *r*(*t* + 1) − *r*(*t*) observed in experiment could be a result of two different ‘random walks’ (blue lines) under reflecting boundary conditions (upper of J). To make a fair comparison with an independent random walk, we sample step sizes Δ*s* of a random walk from a distribution *p*(Δ*s*) (lower left of F) that produces similar shift distribution as in experiment *p*(Δ*r*) (lower right panel of F). (K) Drifts of RFs show distance-dependent correlations, quantified by the average Pearson correlation coefficients. The model can recapitulate the behavior observed in experiment. Shades represent the standard deviation of different pairs of centroids. Experimental results in F-K are plotted using data from (Gonzalez et al. 2019). Parameters: (A)-(C) *β*_1_ = 0.1, *β*_2_ = 0.05, *η* = 0.02, *σ* = 0.01, *N_p_* = 400. (D)-(I): *N_p_* = 200, *α* = 15, *σ* = 0, *η* = 0.01, *β*_1_ = 0, *β*_2_ = 0.05.

As in the simple ring model in the previous section, we found that the drifts of individual RFs can be largely captured by a random walk but with intermittence due to the inactive periods of RFs. The drift speed can be quantified by an effective diffusion constant *D*. The dependence of *D* on the number of neuron *N* is similar to the ring model in the previous section. When *N* is small, sampling noise dominates the synaptic update, resulting in a slight decrease of *D* as *N* increases. However, beyond a certain *N*, *D* increases rapidly with *N* (SI Appendix, Fig. S2E). Our model also predicts that neurons whose RFs have stronger tuning (larger peak amplitude of the RF) tend to be active more often, and have smaller drift (SI Appendix,Fig. S2F,G).

While the above predictions could be compared with long-term recording experiments for animals in a 2D environment, existing long-term recording experiments are limited to 1D environments (typically linear tracks). To compare our model with these experimental results, we simulated our model in a 1D environment, where grid cell responses are modeled as 1D slices through the 2D grid fields, as observed in experiments (Yoon et al. 2016)(Fig. 5D, Material and Methods). The model generates qualitatively similar results as the above 2D place cell model. The learned place fields tile the linear track but drift over time due to ongoing noisy weight updates, yet the representational similarity is stable over time (Fig. 5E). This is also observed in an experiment (Gonzalez et al. 2019), where CA1 pyramidal cells were recorded when mice were in the same familiar environment for several months (Fig. 5F). Due to drifting place fields, the autocorrelation coefficients of neural population vectors in both our model and in experiment decay over time (Fig. 5G). The shift of centroids of place fields increases with time, with a distribution eventually approaching the case wherein the place fields are randomly permuted. Such behavior closely resembles that of experiments (Gonzalez et al. 2019) (Fig. 5H). Despite the continuous reconfiguration of the neural assemblies representing the position, the fraction of active place cells is stable over time (Fig. 5I).

To further explore the underlying structure of centroid shifts, and test the main prediction of our model that the drift of RFs is coordinated, we set out to compare the experiment and our simulation results to a null hypothesis — the shift of RFs behave like an independent random walk. To make a fair comparison, for the null hypothesis, we assume each centroid takes a step size Δ*s* that is drawn from a distribution *p*(Δ*s*) with a reflecting boundary condition (upper panel of Fig. 5J). The distribution *p*(Δ*s*) was chosen such that the resulting centroid shift Δ*r* closely matches that of experiment (lower left panel of Fig. 5J, Material and Methods). Centroid shifts in experiment show clear distance-dependent correlations, i.e., two RFs that are very close to each other are more likely to drift in the same direction on the next day, while RFs that are far apart are more likely to drift in opposite directions (blue line, Fig. 5 K). This is in stark contrast with the independent random walk picture (gray horizontal line, Fig. 5 K), but can be recapitulated by our model (red line, Fig. 5 K), suggesting that the drift of RFs in experiment is coordinated at the population level, possibly to preserve representational similarity.

### A Hebbian/anti-Hebbian network model for drifting RFs of neurons in PPC

The above model and results can be extended to another sensorimotor task, in which mice were trained to navigate a virtual T-maze (Driscoll et al. 2017). At the first half of the T-stem, mice saw one of two alternative visual scenes, and associated them with left-turn or right-turn at the T-junction to receive a reward at the end of the track (Fig. 6A). PPC is essential for this task (Driscoll et al. 2017; Harvey, Coen, and Tank 2012). After learning, a sub-population of neurons in PPC have localized receptive fields, i.e., they fire when a mouse is at a specific position along the T-maze and their RFs tile the T-maze, providing essential information about the task. While mice stably perform the task after learning, the neural population activities in PPC continuously drift over weeks. Despite such drift, the task information can be stably encoded by the activities of a subpopulation of PPC neurons (Driscoll et al. 2017).

**Figure 6:**
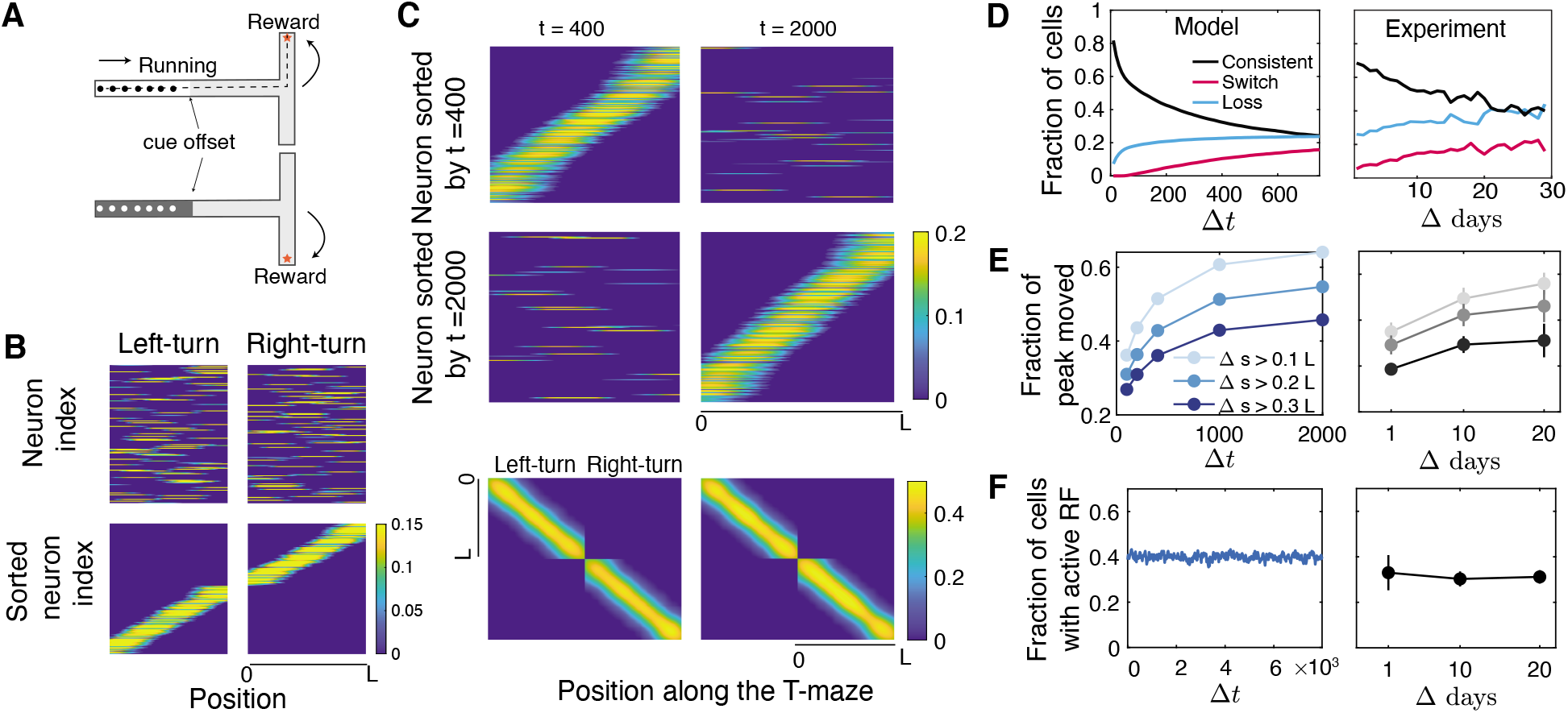
Representational drift in PPC. (A) Schematic of the visual-cue-guided T-maze sensorimotor task as in (Driscoll et al. 2017). The linear length of the track from the beginning to the end (dashed line) is *L*. (B) Population activity for the left-turn and right-turn task before (upper) and after (lower) sorting based on the centroids of their RFs. Only neurons that have active RFs at the given time point are shown. Population activity drifts but representational similarity is stable over time. Activity of neurons identified with significant peak in the sorted time (upper and middle). Representational similarity matrix is stable for both left-turn and right-turn task (lower panels). (D) left: For a group of neurons that have tuning to left-turn (or right-turn) tasks, the fraction of them that have consistent tuning (black), switched tuning (magenta), losing tuning (cyan) to left (or right) in the following time. (E) Shift of RFs for neurons with a significant peak between time *t* and *t* + Δ*t*. Smaller shift happens more often than larger shift. (F) The fraction of the neurons with active RFs is stable across time. In (D)-(F) Left panels are simulation results of our model, right panels are corresponding experiment results from (Driscoll et al. 2017). Parameters: *N* = 400, *α*^2^ = 1.6, *η* = 0.05, *σ* = 10*^−4^, β*_1_ = 10*^−4^, β*_2_ = 10^−3^.

To better understand the positional tuning and drifting behavior of neurons in PPC, we again use a Hebbian/anti-Hebbian network with noisy weight update rule to model this system. For simplicity, the task information is represented by a vector **x***_R/L_*(*θ*) = [cos(*θ*), sin(*θ*), ±1]^┬^, *θ* ∈ [0, *π*), with the last entry indicating a right-turn (1) or a left-turn task (−1).

After learning, the population of output neurons in the model develop positional tuning of the T-maze, i.e., for either left-turn input **x***_L_* or right-turn input **x***_R_*, there are a subpopulation of neurons that fire most strongly when the animal is at the specific positions of the track, forming RFs that tile the maze (Fig. 6B).

To see how the RFs of neurons evolve over time, we first sort neurons with significant RFs based on the centroid positions of their RFs at a reference time point. We find that RFs of neurons drift over time, i.e., neurons rarely have the same or similar RFs at two long-separated time points. However, the population representation of location-context information is stable across time. Thus, at any given time, we can identify a subset of neurons with significant RFs that tile the positions of the T-maze for both left-turn and right-turn tasks (Fig. 6C, upper and middle panels). Despite the drift, representational similarity of both left-turn and right-turn tasks are stable over time (Fig. 6C, lower panel). Neurons also gradually change their tuning to tasks choices. For example, a group of neurons that are tuned to the left-turn tasks at time 0 may loss such tuning or become tuned to right-turn tasks, and vise versa. Drift of an RF accumulates over time, such that the probability of centroid shift that is larger than a certain distance increases with time (left, Fig. 6E). Overall, the fraction of neurons that have positional tuning at any time for both left-turn and right-turn trials are constant (left, Fig. 6F). All these behavior are consistent with the experiment (right panels of Fig. 6D-F). Together, these comparisons shows that our simple model can explain many characteristics of representational drift in PPC.

## Summary and Discussion

In this paper, we explored the hypothesis that representational drift is due to the existence of many (possibly infinite) ensembles of population codes that achieve a representational objective. Noise in learning drives the network to explore this space, causing the drift of population activity. While our focus was on synaptic noise, other sources of noise can also cause similar representational drift with potentially different statistics. Similarly, network architectures that optimize other objective functions can also show drift when learning with noise. However, we expect the drift to be strongly affected by the degeneracy of the solution space of the objective function. For example, in a feedforward network performing online principal component analysis, which has no degeneracy as the principal subspace projection task, we found stabilized representations in the presence of noise (Fig. S3, SI Appendix).

To explore the consequences of our hypothesis in a concrete model, we focused on a well-studied model for biologically plausible representation learning that optimizes similarity-based representational objectives (Pehlevan and Chklovskii 2019). We showed that simple Hebbian/anti-Hebbian networks with noisy synaptic updates recapitulate observed representational drift phenomena in experiments. In the case that the network consists of a single output neuron, we observed that its RF behaves like a random walk on the data manifold with a diffusion constant that depends on the noise amplitude, learning rate, and statistical structure of the input. When the network consists of many neurons, different drifting RFs are coordinated such that representational similarity is stable across time.

We used the Hebbian/anti-Hebbian network as a simplified model to study the RFs of hippocampal place cells and neurons in the PPC. Our model recapitulates the drift statistics at population-level observed in these regions: First, a constant fraction of active neurons represent task variables at a given day. Second, neurons drop in and out of this assembly over days. As a consequence, the autocorrelation coefficient of population vectors decay over time. Third, drift at population level preserves representational similarity (Fig. 5,6). While simple, the network captures the essential properties of those neural circuits, i.e., RFs are shaped by input from upstream and effective lateral inhibition/competition within the layer. It is also possible to model these systems by training a general recurrent neural network (RNN), as has been demonstrated in (Rajan, Harvey, and Tank 2016). It will be interesting to see whether neurons in such RNN models with noisy weight update also show representational drift.

Our model makes several testable predictions. First, our model predicts that the drifts of RFs are coordinated. This coordination is arising from the existence of a representational objective for the neural population as a whole. We verified this prediction in hippocampal data Fig. 5J,K. Second, it predicts that neurons whose synapses have faster turnover dynamics tend to drift more rapidly. For example, the lifetime of spines of pyramidal cells in hippocampus is about 1 to 2 weeks, much shorter than that of neocortex neurons (Attardo, Fitzgerald, and Schnitzer 2015). This suggests that representational drift should be more prominent in hippocampus than in neocortex. Furthermore, the lifetime of synapses can be perturbed by blocking receptors such as NMDA (Zuo et al. 2005), which will alter the stability of RFs. A definitive examination of this prediction requires experiments that both measure the life time of synapses and the long-term neural activity in brain regions that represent learned stereotyped behavior under unperturbed and perturbed states. While challenging, this is nonetheless becoming within reach with new experimental techniques. Third, our model predicts that neurons with strongly tuned RFs should be more stable. This prediction can be tested by examining the amplitude of tuning curves (RFs) of individual neurons and their stability in long-term recording experiments. Furthermore, the strength of RFs can be perturbed by optogenetic tools to examine how it affects the stability of RFs.

Representational drift contradicts the hypothesis that stable neural activity is the substrate of stable behavior. However, there needs to be stable aspects of representations which provide a substrate for stable downstream decoding and readout. Representational similarity can be one such substrate for multiple reasons. First, our modeling shows that achieving stable representational similarity despite the drift of population activity is biologically plausible. Second, stable representational similarity may be a general internal structure of drifting neural population activity. For example, mouse visual cortices show strong representational drift yet the relation between population activities that represent different inputs remains stable and stereotyped (Deitch, Rubin, and Y. Ziv 2021). Conserved and stable internal structure of neural activity has also been discovered in hippocampus and prefrontal cortex in free-behaving mice (Rubin et al. 2019). Third, experimental evidence is consistent with stable representational similarity being a foundation for robust downstream decoding. Studies in monkey motor cortices have shown that stable geometry of latent population dynamics underlies stereotyped reaching tasks (Gallego et al. 2020) despite the inherently variable single neuron activities (Liberti et al. 2016; Rokni et al. 2007) (see however (Chestek et al. 2007; Katlowitz, Picardo, and Long 2018)). Interestingly, a recent experiment has shown that the spatial code of different environments in the hippocampus are random in individual rodents but share the same geometry across different animals (Kinsky et al. 2018). Finally, preserving pairwise similarity of representations may provide some computational benefits. Recent unsupervised learning algorithms for image recognition, such as contrastive representational learning (Chen et al. 2020) and “Barlow Twins” (Zbontar et al. 2021), are based on objectives that maximize representational similarity between a sample and its distorted/augmented versions. Such algorithms can achieve comparable performance to supervised learning algorithms. From a theoretical point of view, the representational similarity matrix (or kernel) determines the number of sampled stimuli required to learn an accurate linear readout from a population code, indicating that performance need not suffer as long as the representational kernel is preserved (Bordelon and Pehlevan 2021).

A hypothesis for achieveing stable readout despite time-varying neural activity is that the variation happens in the “coding-null space” (Druckmann and Chklovskii 2012; Kaufman et al. 2014). Representations in our model exhibit drift in all dimensions, precluding the existence of such coding-null space. Similarly, a closer scrutiny of the response of PPC neurons in the ‘T-maze’ task showed that drift is not confined to a “coding null space” (Rule, Loback, et al. 2020). Hence, an adaptive readout mechanism which involves synaptic plasticity to track and compensate the drift is required to achieve stable behavior (Rule, Loback, et al. 2020; Rule and O’Leary 2021). Whether and how such a mechanism is implemented in the brain remains an open question.

The ubiquity of representational drift raises the question of whether it is a biological feature or a bug. Representational drift may be desirable under certain circumstances (Mau, Hasselmo, and Cai 2020). For example, in a model of the bird song learning system, variation in the neural representation of the stereotyped behavior enables the system to adapt quickly to a shift of target song, and to reduce error due to loss of neurons (Duffy et al. 2019). Drift can accommodate new learning with minimal inference by continuously modifying existing memories (Mau, Hasselmo, and Cai 2020). Other authors proposed that noisy synaptic plasticity and spine motility enable cortical networks of neurons to carry out probabilistic inference by sampling from a posterior distribution of network configurations (Kappel et al. 2015). Such sampling would lead to a representational drift as a byproduct.

## Material and Methods

### Similarity matching and the linear Hebbian/anti-Hebbian network

The linear Hebbian/anti-Hebbian network can be derived from (1). The detailed derivation can be found in (Pehlevan, Hu, and Chklovskii 2015; Pehlevan, Sengupta, and Chklovskii 2018), we sketch the main steps here. Starting from the cross term in (1), by introducing a new matrix variable **W** ∈ ℝ^*k*×*n*^, we obtain

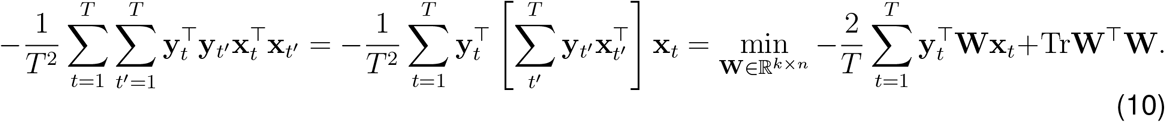

Similarly, we can introduce another matrix variable **M** for the quartic **y***_t_* term in (1):

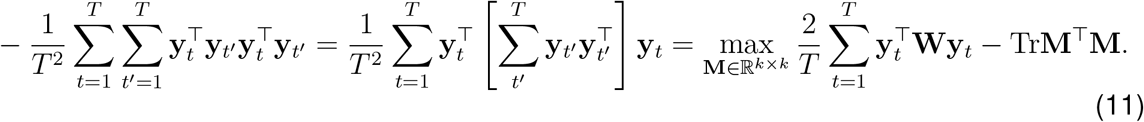

By substituting (10) and (11) into (1) and changing orders of optimization (Pehlevan, Sengupta, and Chklovskii 2018) we get:

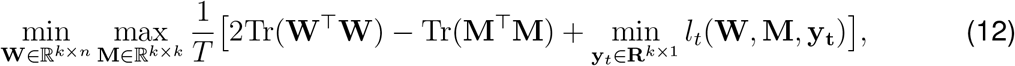

where

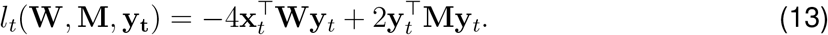

The minimax problem (12) can be solved by the following two-step online algorithm. First, minimizing (13) while keeping **W** and **M** fixed, which is solved by running the dynamics of output variable **y***_t_* until convergence

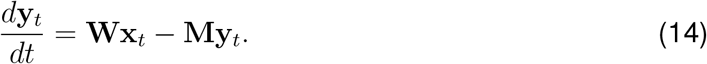

Second, after the convergencec of **y***_t_*, update **W** and **M** by gradient descent and gradient ascent of (12) respectively:

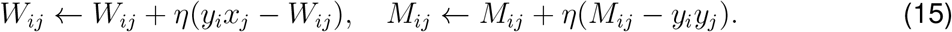

The above learning algorithm (14), (15) can be naturally mapped onto a single-layer biologically plausible neural network, the linear Hebbian/anti-Hebbian network. Here, **y***_t_* is the neural activity of the output, **W** and **M** are synaptic matrices of the forward and lateral connections respectively. The synaptic update rule (15) is local since the change of a synapse only depends on the activity of presynaptic and postsynaptic neurons.

### Calculation of the rotational diffusion constant

An analytical calculation of the rotational diffusion constant, defined by (Hunter et al. 2011; Kämmerer, Kob, and Schilling 1997; Mazza et al. 2006),

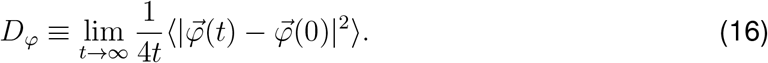

is difficult. However, we were able to obtain an approximation which matches numerical experiments very well, as shown in Fig. 2E-G. We present the details of this derivation in the SI Appendix. Our approximation assumes that 1) angular displacements of the representation vectors after different time steps are not correlated, and 2) the network weights stay close to the optimal representation manifold. Under these assumptions, *D_φ_* can be approximated by the mean squared angular displacement (MSAD),

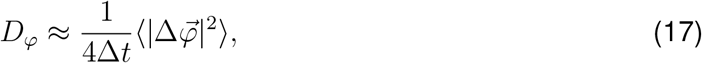

where Δ*t* is the small time interval elapsed during a single step update, and 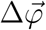 arises from a noisy synaptic update to the network with an optimal set of synapses. We calculate MSAD analytically (SI) to arrive at (5).

To numerically estimate *D_φ_* from trajectory of **y**(*t*) with total length of *T* time steps, we first calculate 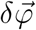 at each simulation step, then estimate 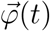 by cumulatively summing 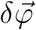 up to time step *t*. Next, we estimate the MSAD of time interval *τ* using all the pairs of 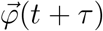 and 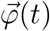, which gives 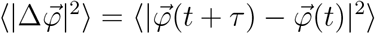. Last, we plot 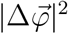 as a function of *τ* and fit a line that pass the origin to the data. The slope of the best fit is then 4*D_φ_*.

### Hebbian/anti-Hebbian network and nonnegative similarity matching

The nonlinear Hebbian/anti-Hebbian network (Eq. s(7) and (8)) can be derived from the general nonnegative similarity matching (NSM) problem (Pehlevan 2019; Pehlevan and Chklovskii 2019). Denoting the input data as a set of vectors **x**_*t*=1,…, *T*_ ∈ ℝ^*n*^ and the corresponding output vectors **y**_*t*=1,…,*T*_ ∈ ℝ^*k*^, the NSM objective is defined as

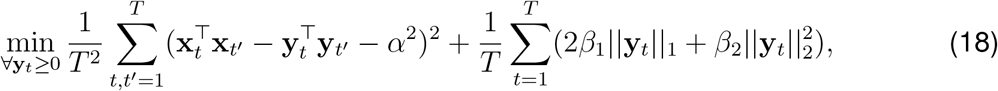

where *α*^2^ sets the threshold of similarity to be preserved in the output representation, the other two regularizers *β*_1_, *β*_2_ control the sparsity and amplitude of output. The detailed derivation of and (8) from (18) is described in (Pehlevan 2019).

To see why the above NSM objective (18) leads to localized RFs, we can consider the simpler case where *β*_1_ = *β*_2_ = 0 and a single pair of inputs. If two inputs are similar, i.e., **x**_1_ · **x**_2_ > *α*^2^, then the corresponding outputs **y**_1_ and **y**_2_ would prefer **y**_1_ · **y**_2_ = **x**_1_ · **x**_2_ − *α*^2^, i.e., they are also similar. In contrast, if two inputs are less similar, i.e., **x**_1_ · **x**_2_ < *α*^2^, due to the nonnegativity of outputs, **y**_1_, **y**_2_ they tend to be orthogonal: **y**_1_ · **y**_2_ = 0. To achieve this, dissimilar inputs must activate non-ovelapping sets of neurons. Thus, in manifold learning, (18) preserve local geometric structure in the **y** representation space of the input data clouds. A detailed explanation of why localized RFs are learned in a simplified version of (18) is provided in (Sengupta et al. 2018).

The neural dynamics derived from (18) (with regularizers) differ from that in main text slightly by changing the transfer function in (7) to

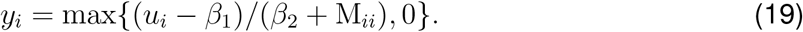

### Derivation of the diffusion constant of the ring model

We sketch the derivation of (9) here, more details are in SI Appendix. We again consider the approximation that the diffusion coefficient can be approximated by the mean squared displacement around a fixed point by a noisy synaptic update.

Consider a single output neuron that learns a RF from inputs that are on a ring manifold (Fig. 3A). The response of the output neuron to **x** = [cos *θ*, sin *θ*]^┬^ is

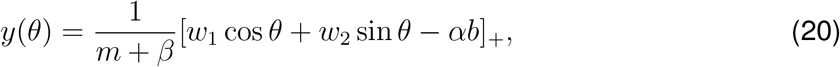

where [*x*]_+_ denotes the rectified linear function and *β* is the *l*_2_ regularizer. The stationary state parameters 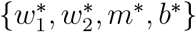 satisfy the following conditions

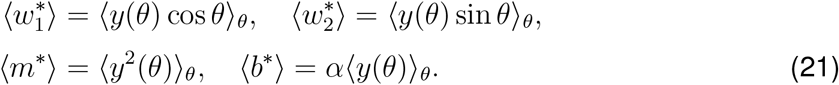

These equations can be solved self-consistently by assuming an ansatz of the form:

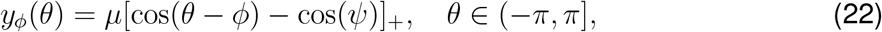

which gives the dependence of *μ* and *ψ* on *α, β*

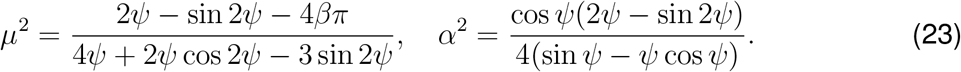

Using the fact that *dy*(*θ*)/*dθ* = 0 at *θ* = *ϕ*, we have 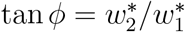 and

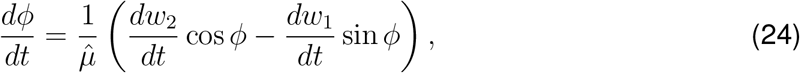

where 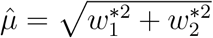 is the norm of weight vector. Using the noisy update rule (8) and (22), the shift of centroid due to one-step update eventually becomes

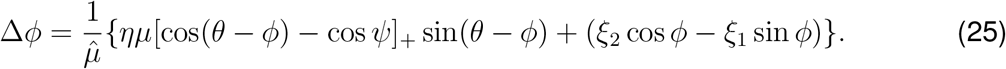

Finally, using the relation 〈(Δ*ϕ*)^2^〉 ≈ 2*D*Δ*t*, we have

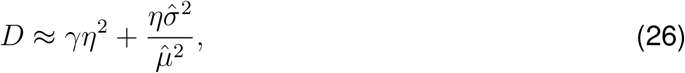

where 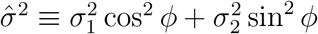 and

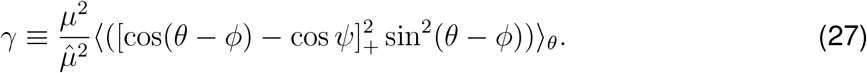

When *α = β* = 0, *σ*_1_ = *σ*_2_ = *σ*, we have *γ* = 1 and 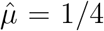, (26) reduces to (9) in the main text.

### Numerical simulation of 2D place cells

We considered a 32 × 32 grid plane as the environment, each position (*x, y*) is represented by a group of grid cells with different grid spacings, orientations and offsets as observed in experiment (Stensola et al. 2012). The hexagonal firing fields of grid cells are modeled as a summation of three two-dimensional sinusoidal functions as in (Kropff and Treves 2008; Lian and Burkitt 2020; Solstad, E. I. Moser, and Einevoll 2006)

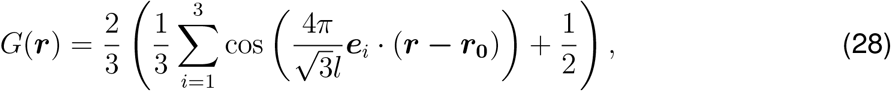

where ***r*** = [*x, y*]^┬^ is the location on the plane, ***r***_0_ = [*x*_0_, *y*_0_]^┬^ is the phase offset, *l* is the grid spacing. ***e****_i_* = (cos(2*πi/*3 + *θ*), sin(2*πi/*3 + *θ*)), *i* = 1, 2, 3 is the unit vector in the direction 2*πi/*3 + *θ* with *θ* being the grid orientation. In the simulation, grid cells have 5 modules, i.e., *N_l_* = 5. The value of *l* increases as geometric series with a ratio 1.42 that is consistent with experiment (Stensola et al. 2012). For example, in our simulation, the smallest spacing is 0.2*L* with *L* being the linear length of the plane, then the rest spacing would be 0.2 × 1.42*L*, …, 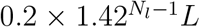. In each module, the number of orientation *θ*, *N_θ_* = 6, which are drawn uniformly in the range [0, *π/*3). Similarly, the number of grid phase offsets *x*_0_, *y*_0_ are *N_x_* = 5 and *N_y_* = 5, which are drawn uniformly in the range [0, *l*). As result, the total number of grid cell is *N_g_* = *N_l_N_θ_N_x_N_y_* = 750.

### Numerical simulation of 1D place cell

We consider a linear track with length *L*. Tuning curves of grid cells on the linear track are slices through 2D grid fields described above. The orientation of the slices are the same and randomly selected in the range [0, *π/*3].

### Autocorrelation coefficient of the population vector

In all the cases, the autocorrelation coefficient *ρ* of population vector is defined as the Pearson’s correlation coefficent between **y***_t_* and **y**_0_ to the same input:

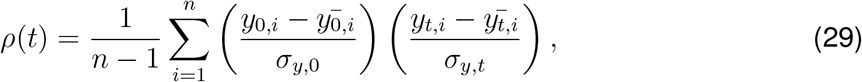

where 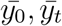 are the mean of *y*_0,*i*_ and *y_t,i_*, *σ*_y,0_, *σ*_*y,t*_ are the standard deviation of *y*_0,*i*_ and *y_t,i_*.

### Step size in independent random walks place fields

In Fig. 5J,K, the step size of independent random walks were drawn from a distribution *p*(Δ*s*) closely matching that of experiment. To determine this distribution, we first calculated the distribution of centroid shift between two adjacent days in experiment *p*(Δ*r*) with Δ*r = r*(*t* + 1) − *r*(*t*). For a random walk whose centroid is at position 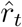 its position at next time step is 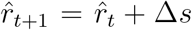 with Δ*s* randomly sampled from *p*(Δ*s*). To constrain 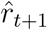 in the range of the track [0, *L*] with *L* being the length of the track, we assumed a reflecting boundary condition, which gives

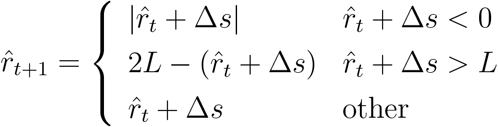

The shift of centroid in the random walk model is then determined by 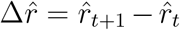 according to the above equation. Our aim is to find a distribution *p*(Δ*s*), such that 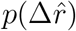 is close to that of experiment *p*(*r*). Based on the shape of *p*(*r*), we searched *p*(Δ*s*) from a family of Levy’s alpha stable distribution (Samorodnitsky and Taqqu 2017) by minimizing the Kullback–Leibler divergence between *p*(*r*) and 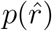

### Data source and processing

Experimental data presented in Fig. 5 are originally described in (Gonzalez et al. 2019). We used the processed data and MATLAB code, which are available at the Caltech Research Data Repository (https://doi.org/10.22002/d1.1229) to produce these plots.

Experimental data presented in Fig. 6 is extracted from Figure 2C, 2D, and 4E of (Driscoll et al. 2017). The data is freely available in (Driscoll et al. 2020).

## Acknowledgements

This work was supported by NIH (1UF1NS111697-01), the Intel Corporation through Intel Neuromorphic Research Community, and a Google Faculty Research Award. We thank Walter Gonzalez, Hanwen Zhang, Anna Harutyunyan and Carlos Lois for sharing the data on place cell recordings. We thank Laura Driscoll, Noah Pettit, Matthias Minderer, Selmaan Chettih and Christopher Harvey for making the T-maze experimental data available. We are grateful to members of Pehlevan group for helpful discussions, and Christopher Harvey for comments on the manuscript.

## Competing interests

The authors declare no competing interests.

## I. DERIVATION OF THE ROTATIONAL DIFFUSION CONSTANT THE IN THE LINEAR HEBBIAN/ANTI-HEBBIAN NETWORK

In this section, we derive an analytical expression for the rotational diffusion constant defined by [1, 2]

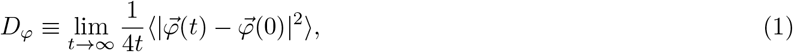

where brackets mean averaging over different realizations of the noise. Obtaining an exact expression for *D_φ_* is difficult, but we are able to derive an approximation that matches numerical experiments well, as shown in Figure 2 E, F and G of main text.

Our approach relies on two simplifications. First, we define the single-step angular displacement

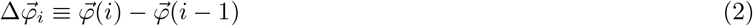

and note that

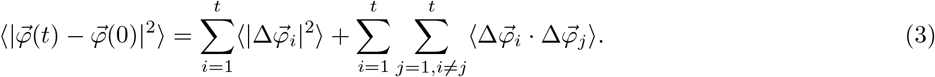

We assume that the correlation between angular displacements at different times is negligible. Therefore, we approximate

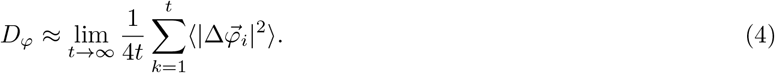

Second, we assume that the network weights start at a configuration that is already an optimal solution to the similarity matching objective, projecting the input to its principal subspace, and the drift keeps the weights in the optimal solution space. This is a reasonable approximation because of a linear stability analysis presented in [3, 4]. We now review that argument. We refer an optimal solution to the similarity matching problem in the offline setting without noise as a fixed point and denote it with a^. We note that a general perturbation of feature map *δ***F** around a fixed point 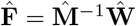 can be decomposed as

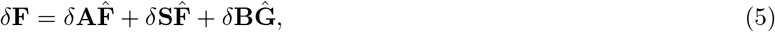

where *δ***A** is a *k* × *k* antisymmetric matrix, *δ***S** is a *k* × *k* symmetric matrix, and 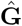 is a (*n* − *k*) × *n* matrix with orthonormal rows. These rows are chosen to be orthogonal to the rows of **F**. *δ***B** is a *k* × (*n* − *k*) matrix [4]. So we have 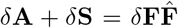. The first term corresponds to a rotation of the neural filter basis of the principal subspace, the second term captures deviations from orthogonality of the basis vectors within the subspace, and the third term captures perturbations of the weight vectors that lead to projecting outside the principal subspace. As shown in [4], the fixed point is stable to the perturbation due to the second and third term, meaning they decay exponentially to zero, making a principal subspace projection linearly stable. Therefore, we consider drift due to the first term, which rotates neural filters and, in turn, the data cloud. We find that (see below) in this limit, 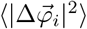 is independent of time step *i*. Therefore, our final approximation is

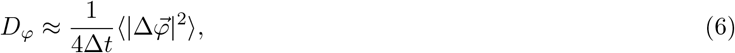

where Δ*t* is the small time interval elapsed during a single step update, and 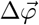 arises from a noisy synaptic update to the network with an optimal set of synapses. This quantity is called mean squared angular displacement (MSAD). This approximation turns out to match simulations very well as shown in Figure 2 E, F and G.

Next, we calculate *D_φ_*. In the linear Hebbian/anti-Hebbian network for principle subspace projection task, the learning rule with synaptic noise is

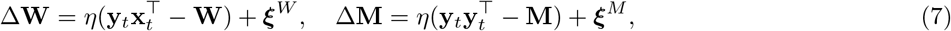

where 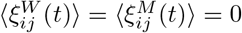 and 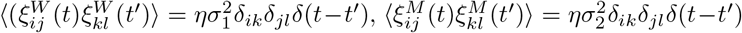. As discussed, by estimating the variance of the rotation the learned representation during a single-step update under rule (7), we can define an effective rotational diffusion constant that is related to this variance. More specifically, in the small update and noise regime, *δ***A** is related to an infinitesimal rotation **R** by 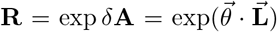 where 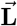 is the infinitesimal rotation generator [5]. 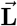 is a tensor, whose components can be written in matrix form. We start by writing *δ***F** in terms of the perturbation of 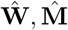

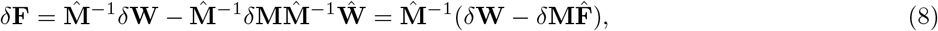

where we have used the property 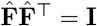 Right-multiplying (8) by 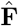 and using (7), we have

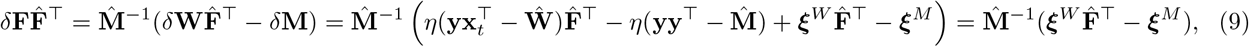

where we have used the fact

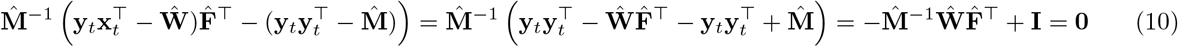

Now, the antisymmetric part 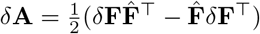 can be written down explicitly:

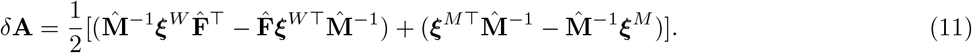

The mean squared angular displacement (MSAD) is related to *δ***A**. To see this more clearly, consider a *d*-dimension rotation, which can be interpreted as rotation in a *d* − 1 dimensional hyperplane from one unit vector to another unit vector. Given any two *d*-dimensional orthogonal unit vector **e_1_**, **e_2_**, i.e., **e_1_ ^┬^ ·e**_1_ = **e_2_ ^┬^ ·e**_2_ = 1, **e_1_ ^┬^ ·e**_2_ = **e_2_ ^┬^ ·e**_1_ = 0. The generator for this rotation can be represented as

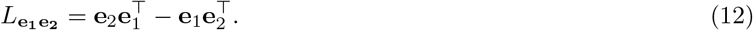

Hence *δ***A** can be expressed as 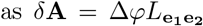, with Δ*φ* reflecting the rotation ‘amplitude’. Using the fact that 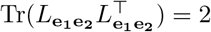, we have

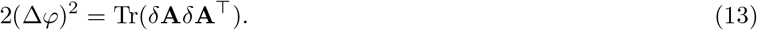

The variance of *δA_ij_* is

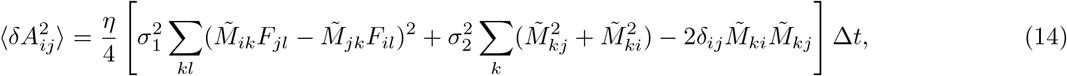

where 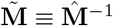 and the average 〈〉 is over the noise distribution Δ*t*, is the time interval of the single-step update.

Using the fact that 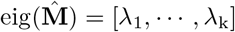 [4], and 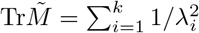 We have

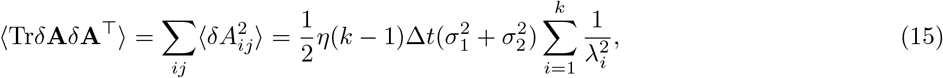

where we have used the fact. We then define the rotational diffusion constant *D_φ_* by the relation 〈|*φ*(*t*+Δ*t*)−*φ*(*t*)|^2^〉 = 2(*k* − 1)*D_φ_*Δ*t*. With Eq.(13) and Eq.(15), we arrive Eq. 5 in the main text.

## II. DERIVATION OF THE EFFECTIVE DIFFUSION CONSTANT IN THE RING MODEL

Here, we calculate the diffusion constant in the ring model for a single output neuron, using the MSAD approximation as before.

We start from the simplest setup for the 1D place cell model: a single place cell which receives input from the “ ring” manifold, i.e., the position is parameterized as **x** = [cos *θ*, sin *θ*]^┬^. The response of the neurons is given by

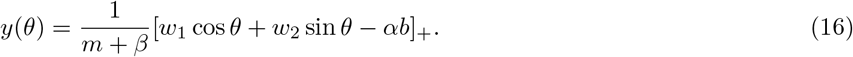

Here and after, we use [*x*]_+_ to denote the rectified linear function. We define a steady state where the average update to the weights is zero. Denoting the stationary state weights as 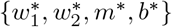 this leads to the conditions:

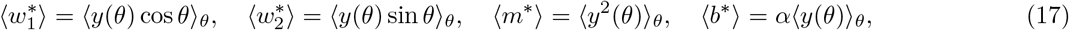

Where 〈·〉_θ_ means averaging over *θ* ∈ [− *π, π*) which is a uniform distribution. These equations can be solved self-consistently using an ansatz of the form:

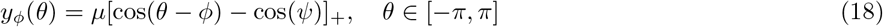

where*ψ* determines the “width” of the RF, and *μ*(1 − cos*ψ*) is the peak amplitude and *ϕ* is the centroid of the receptive field. Plugging (18) into (16) and (17), we find that

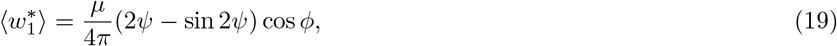

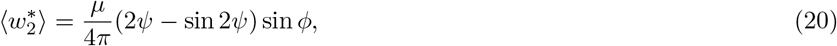

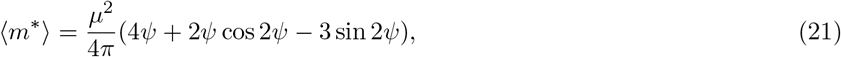

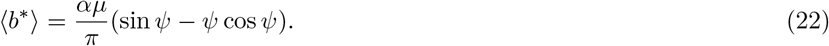

(16) can be rewritten as Compared with (18), we have

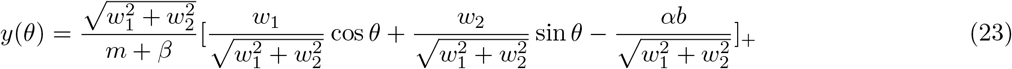

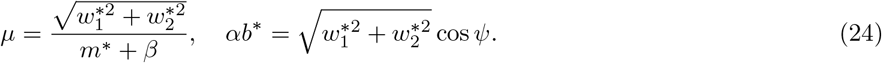

Combining (19)–(22) and (24), we get the dependence of *μ* and on *α, β*, given parametrically by

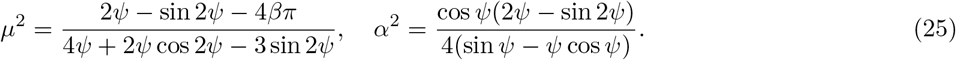

Next, we proceed to estimate the drift due to noisy synaptic updates. From (23), we have

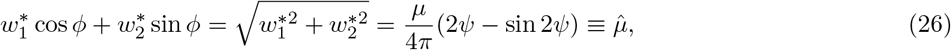

where we have defined 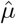 to simplify the following notations. Using the fact that *dy*(*θ*)*/dθ* = 0 at *θ* = *ϕ*, we have 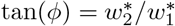 and

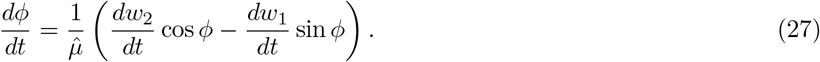

We are interested in how the centroid of the RF changes when a perturbation is added to the stationary weight vector

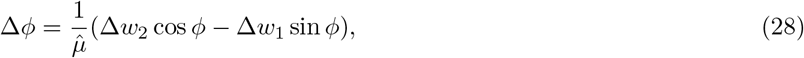

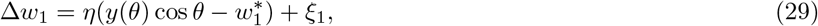

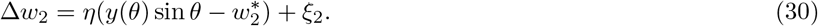

The Gaussian white noise terms have the following property: 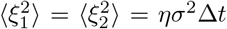 with Δ*t* as the time interval between two adjacent update events. From (26), we have 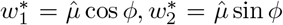 Then Δ*ϕ* can be written as

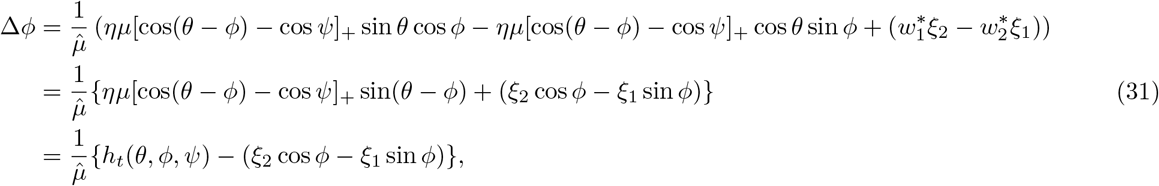

where we have defined *h_t_*(*θ, ϕ, ψ*) = *ημ*[cos(*θ* − *ϕ*) − cos *ψ*]_+_. Since in the online learning, *θ* is sampled randomly, we can regard *h_t_* as a stochastic process. Averaging over *θ*, we have

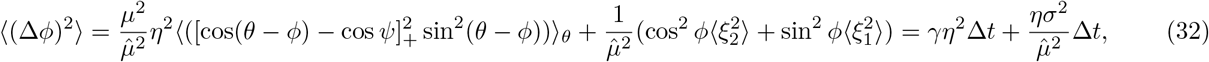

where

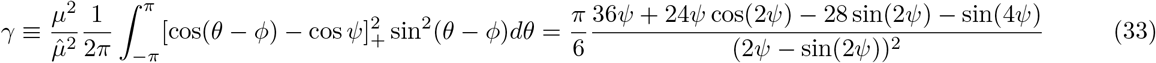

Using the relation 〈(Δ*ϕ*)^2^(Δ*t*)〉 = 2*D*Δ*t*, we have

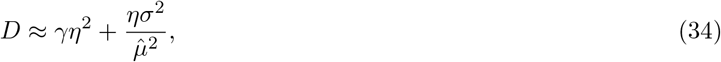

where we use ≈ because we calculate single-step MSAD.

---

**FIG. S1.**
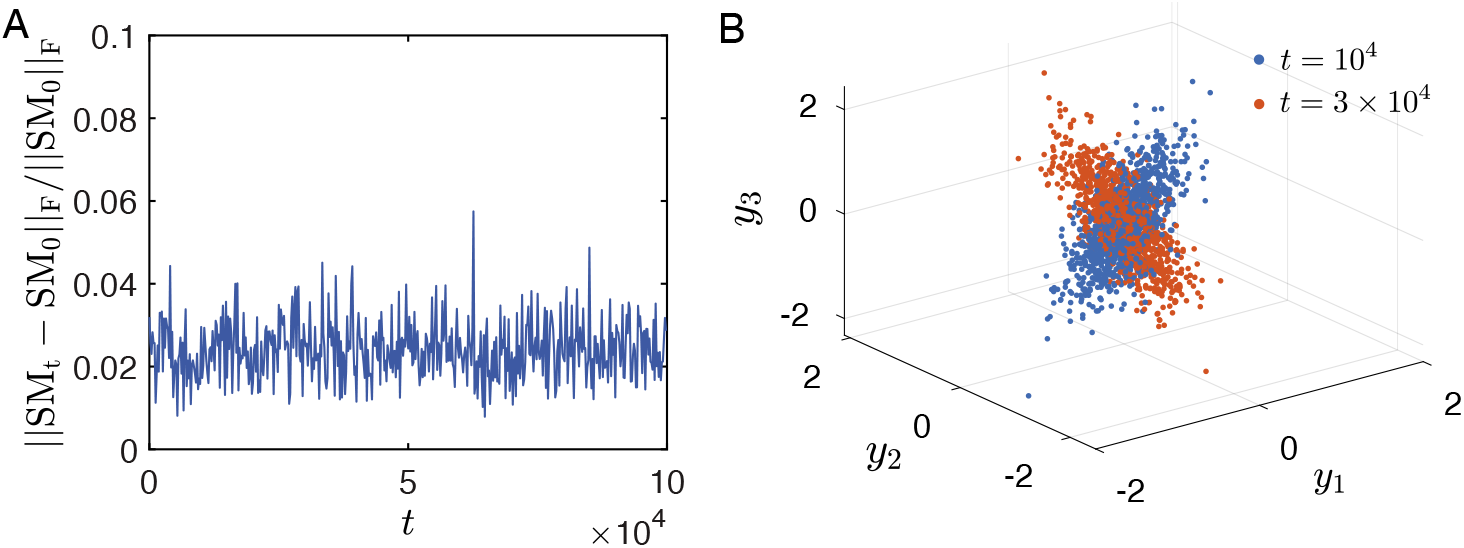
(A) The relative change of Frobenus norm of the similarity matrix at time *t* compared with time point 0 in the PSP task. (B) Ensemble of output **Y** ≡ [**y**_1_, …, **y**_1_] at two time points. The data clouds have ellipsoid shape. Related to figure 2 in the main text. Parameters are the same as in figure 2 of the main text.

**FIG. S2.**
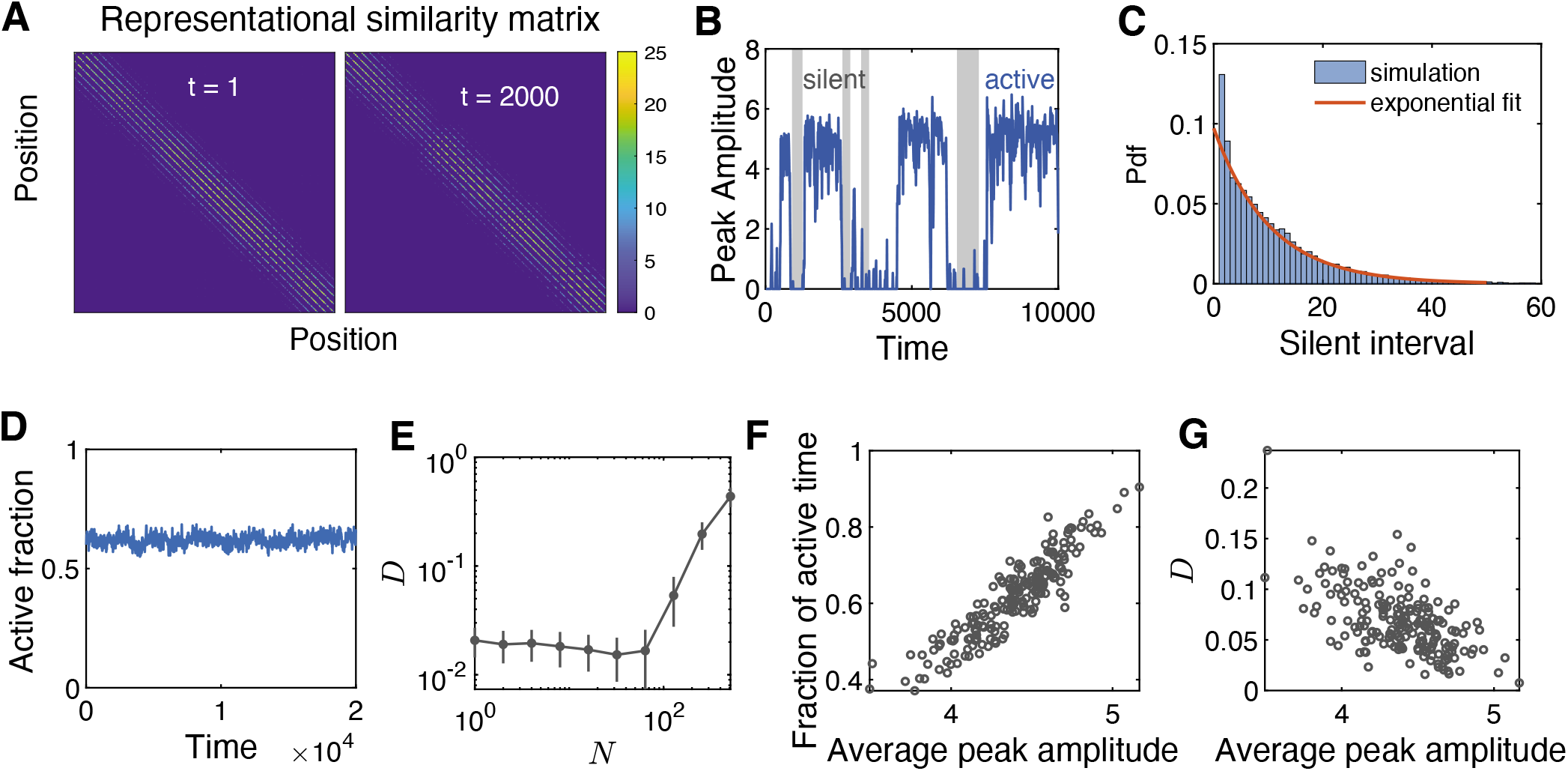
Drift of 2D place cells in the model. (A) Representational similarity is preserved despite the continuous drift of place cell RFs. Positions on the plane are represented by an index from 1 to 1024. (B) The RFs are intermittent. The peak amplitude of an example place field has active and silent bouts. (C) The interval of silent bouts follow exponential distribution. At population level, there is a constant fraction of active RFs over time. (E) Dependence of effective diffusion constant on the total number output neurons. (F,G) Place cells that have stronger place fields tend to be active more often (F) and also more stable as quantified by smaller diffusion constant (G). Parameters used are the same as Figure 5 in the main text.

**FIG. S3.**
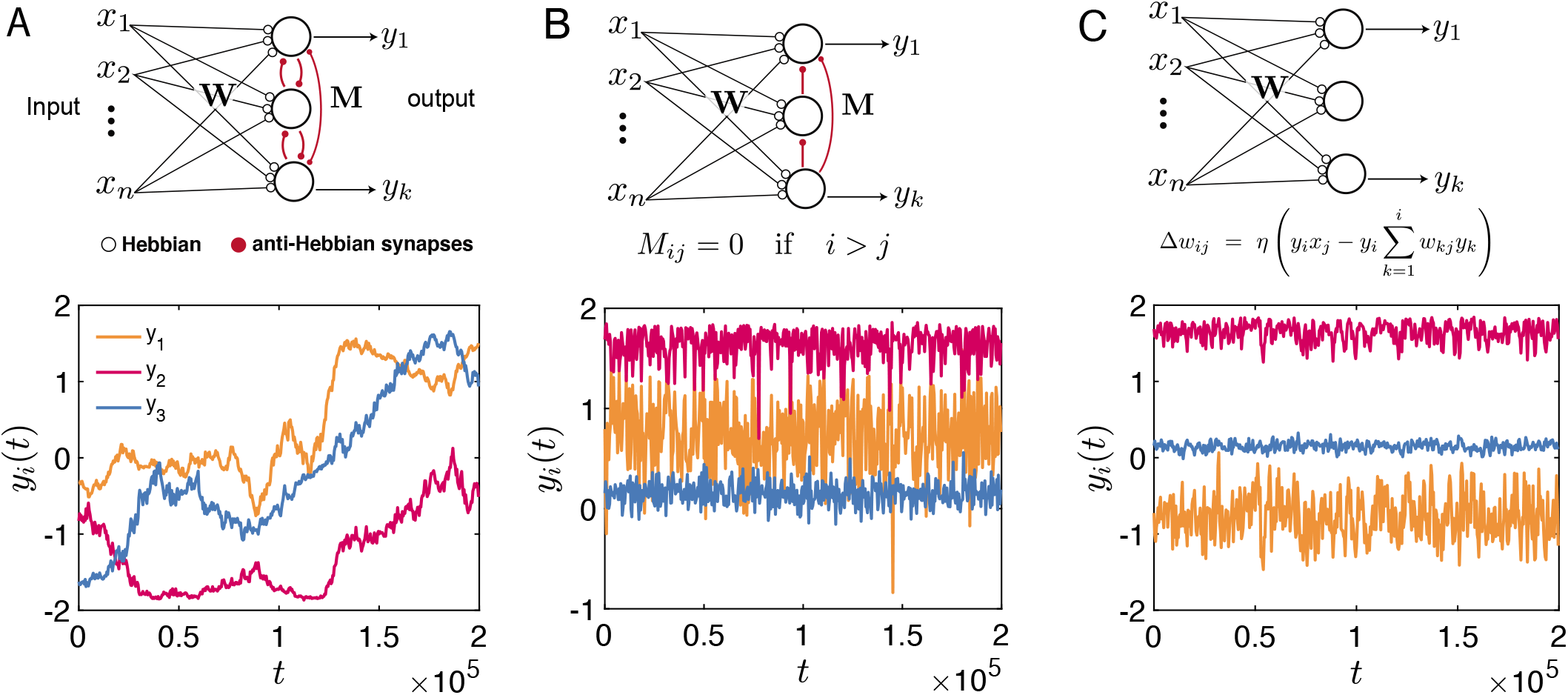
Degeneracy of learning objective function and representational drift. We compare the long-term behavior of learned representations in three different networks. (A) Upper: the Hebbian/anti-Hebbian network for PSP. Lower: the evolution of the three components of a representation **y***t*. (B) Upper: The network differs from Hebbian/anti-hebbain network only in the lateral matrix **M** which break the rotational symmetry of PSP solution. The learning rule is the same. Lower: the learned representation only fluctuates around a equilibrium. (C) A single feedforward network that perform online principle component analysis with Sanger’s rule [6]. This network has only feedforward input matrix **W** and the learning rule is nonlocal. Lower: learned representation is relatively stable in the presence of noise. Parameters are the same as in the figure 2 of main text except that *η* = 0.01.

## Notes

### Competing Interest Statement

The authors have declared no competing interest.

